# Transcription of HIV-1 at sites of intact latent provirus integration

**DOI:** 10.1101/2024.04.26.591331

**Authors:** Ana Rafaela Teixeira, Cintia Bittar, Gabriela S. Silva Santos, Thiago Y. Oliveira, Amy S. Huang, Noemi Linden, Isabella A.T.M. Ferreira, Tetyana Murdza, Frauke Muecksch, R. Brad Jones, Marina Caskey, Mila Jankovic, Michel C. Nussenzweig

## Abstract

HIV-1 anti-retroviral therapy is highly effective but fails to eliminate a reservoir of latent proviruses leading to a requirement for life-long treatment. How the site of integration of authentic intact latent proviruses might impact their own or neighboring gene expression or reservoir dynamics is poorly understood. Here we report on proviral and neighboring gene transcription at sites of intact latent HIV-1 integration in cultured T cells obtained directly from people living with HIV, as well as engineered primary T cells and cell lines. Proviral gene expression was correlated to the level of endogenous gene expression under resting but not activated conditions. Notably, latent proviral promoters were 100-10,000X less active than in productively infected cells and had little or no measurable impact on neighboring gene expression under resting or activated conditions. Thus, the site of integration has a dominant effect on the transcriptional activity of intact HIV-1 proviruses in the latent reservoir thereby influencing cytopathic effects and proviral immune evasion.

## Introduction

HIV-1 proviral integration into the genome of CD4^+^ T cells typically leads to production of up to 10^4^ virions per cell and results in cell death by apoptosis (Wei et al., 1995; Perelson et al., 1996). However, in a small number of instances the integrated provirus remains latent but can be reactivated *in vitro* or *in vivo* upon treatment interruption (Pace et al., 2011; Cohn et al., 2020; Liu et al., 2020; Dufour et al., 2021; Siliciano and Siliciano, 2022). Anti-retroviral therapy (ART) prevents the spread of productive infection and disease progression but does not eliminate a reservoir of infected cells that carry intact latent proviruses (Wong et al., 1997; Chun et al., 1998; Finzi et al., 1999). This reservoir represents the primary barrier to HIV-1 cure (Li et al., 2016; Margolis and Archin, 2017).

Obtaining a detailed understanding of the reservoir has been challenging because the cells that carry intact latent proviruses are rare and there are no definitive cell surface markers that can be used to isolate them (Ho et al., 2013; Crooks et al., 2015; Bachmann et al., 2019; Darcis et al., 2019; Cohn et al., 2020; Peluso et al., 2020; Weymar et al., 2022; Sun et al., 2023; Wong et al., 2023). Nevertheless, there has been a great deal of progress in understanding the nature of the reservoir (Pace et al., 2011; Cohn et al., 2020; Liu et al., 2020; Dufour et al., 2021; Siliciano and Siliciano, 2022). Documented features of the reservoir include: 1. CD4^+^ T cells that carry intact latent proviruses are oligoclonal (Lorenzi et al., 2016; Simonetti et al., 2016; Bui et al., 2017; Hosmane et al., 2017; Cohn et al., 2018; Lu et al., 2018); 2. the clones of cells carrying intact latent proviruses are dynamic and while the overall number of T cells carrying latent proviruses decreases over time they also become more clonal (Cho et al., 2022); 3. intact proviruses found in large CD4^+^ T cell clones are the least likely to be reactivated (Lorenzi et al., 2016) 4. some intact proviruses in the latent reservoir are transcriptionally active, and others are not (Einkauf et al., 2022).

During initial infection, integrase favors proviral deposition in the introns of highly expressed genes associated with regions of accessible chromatin (Schröder et al., 2002; Han et al., 2004; Craigie and Bushman, 2012). These proviruses are the first to be eliminated due to their cytopathic effects. Over time on ART, and in elite controllers, intact proviruses are enriched in heterochromatin, non-genic regions, and in an opposite orientation to the host gene (Einkauf et al., 2019; Jiang et al., 2020; Huang et al., 2021; Einkauf et al., 2022). In addition to centromeric and satellite DNA, integration into zinc finger (ZNF) genes is enriched possibly because these sites are less permissive for provirus expression and subsequent negative selection (Jiang et al., 2020; Huang et al., 2021). Consistent with this idea experiments in transformed cell lines using randomly integrated HIV-1 reporter proviruses suggest that transcriptional activity is dependent on the site of integration (Jordan et al., 2001; Lewinski et al., 2005; Sherrill-Mix et al., 2013; Chen et al., 2017; Battivelli et al., 2018; Collora and Ho, 2023). However, the relationship between the site of authentic latent proviral integration, and transcriptional activity or its effect on neighboring genes is not known. Understanding how the reservoir of intact latent proviruses is regulated and how it impacts neighboring gene expression in primary CD4^+^ T cells is critical to understanding how the reservoir is maintained and how it might be eliminated.

To address these questions, we analyzed HIV-1 and global gene expression in Jurkat and primary T cells that carry HIV-1 reporters inserted precisely into sites of intact latent provirus integration and in authentic latent CD4^+^ T cells isolated directly from people living with HIV (PLWH).

## Results

### HIV-1 LTR reporter expression by flow cytometry

To determine how intact proviruses alter gene expression or are influenced by their position in the host genome we selected 7 well documented sites of latent HIV-1 integration obtained from PLWH, 3 of which had been demonstrated to be inducible *in vitro* (Fig.1 A) (Jiang et al., 2020; Huang et al., 2021; Einkauf et al., 2022). The 7 sites differed in their documented level of host gene expression, proviral orientation with respect to the neighboring gene, and whether HIV-1 transcripts were detected in primary cells (Fig. 1 A). Five of the 7 sites were found among expanded clones and 2 in unique CD4^+^ T cells (Jiang et al., 2020; Huang et al., 2021; Einkauf et al., 2022). One was in a non-genic region of chr13, and the remainder were in introns of genes expressed at low (*ZNF407, ZNF140*), medium (*KDM2A, ZNF460*, *KCNA3*) and high (*ATP2B4*) levels (Fig. 1 A).

**Figure 1.**
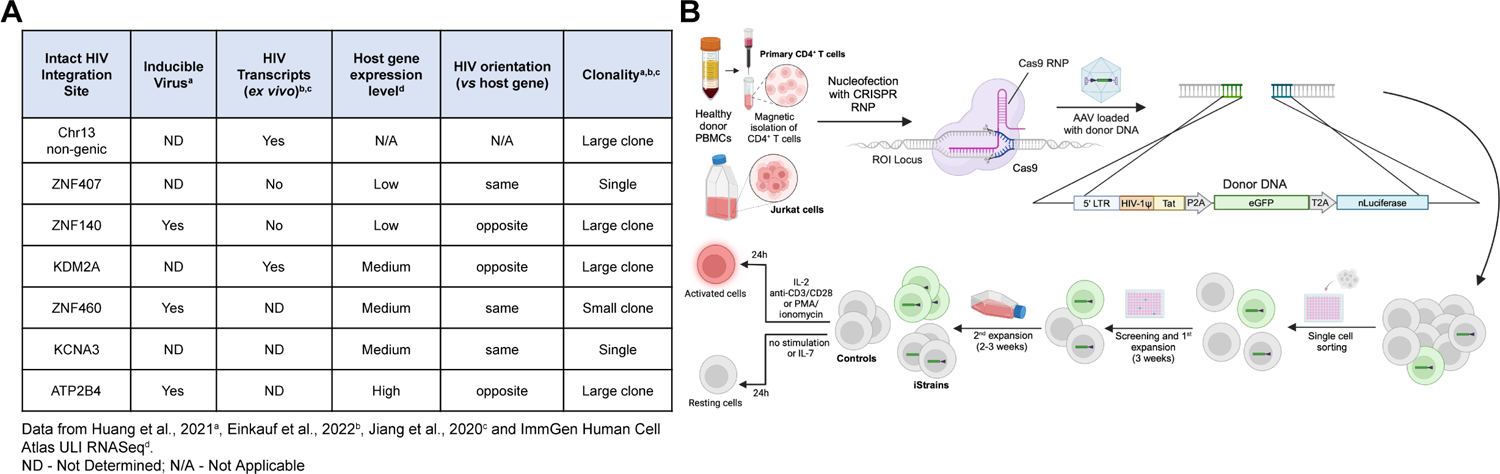
Selected integration sites and production of reporter T cell clones. **(A)** Table of selected integration sites. **(B)** Schematic representation of the methods used to produce reporter T cell lines (see Materials and methods section for details). Created with BioRender.com.

A reporter construct consisting of an HIV-1 Long Terminal Repeat (LTR), transactivator of transcription (Tat), green fluorescent protein (eGFP) and nano Luciferase (nLuciferase) was inserted into the precise position of each of the 7 intact latent proviruses using CRISPR ribonucleoproteins (crRNPs) in Jurkat cells or primary CD4^+^ T cells. Integration was verified by PCR in expanded clones obtained from single cells (iStrains; Fig. 1 B). The amplified region encompassed the integration site and construct, and the integrity of both was confirmed by Sanger sequencing. Several independent clones were obtained for *ATPB2B4* and *KCNA3* in Jurkat and CD4^+^ T cells, and for *ZNF407* in Jurkat cells. We were unable to obtain an integration in *ZNF407* in primary CD4^+^ T cells. Control cells were transfected and cultured under the same conditions but did not contain a reporter.

As measured by flow cytometry, reporter expression varied depending on the integration site. There was no detectable expression by reporters integrated into *ZNF140*, and *ZNF407,* or a non-genic region in Chr13 in Jurkat cells or primary CD4^+^ T cells under resting conditions or after activation with phorbol 12-myristate 13-acetate (PMA) and Ionomycin or anti-CD3 +-CD28 monoclonal antibodies, respectively (Fig. 2 A and 3 A). The reporters integrated into *ZNF460* and *KDM2A* were also silent in primary CD4^+^ T cells but in contrast to the other 3 silent genes, they were expressed in a fraction of the cells upon activation (Fig. 3 A). Reporters integrated into *KCNA3* and *ATP2B4* from multiple independent clones were expressed under resting conditions and expression was further enhanced by activation, as measured by an increase in the percentage of cells expressing GFP and in their mean fluorescence intensity (Fig. 2 A, 3 A and Fig. S1 A and S1 B).

**Figure 2.**
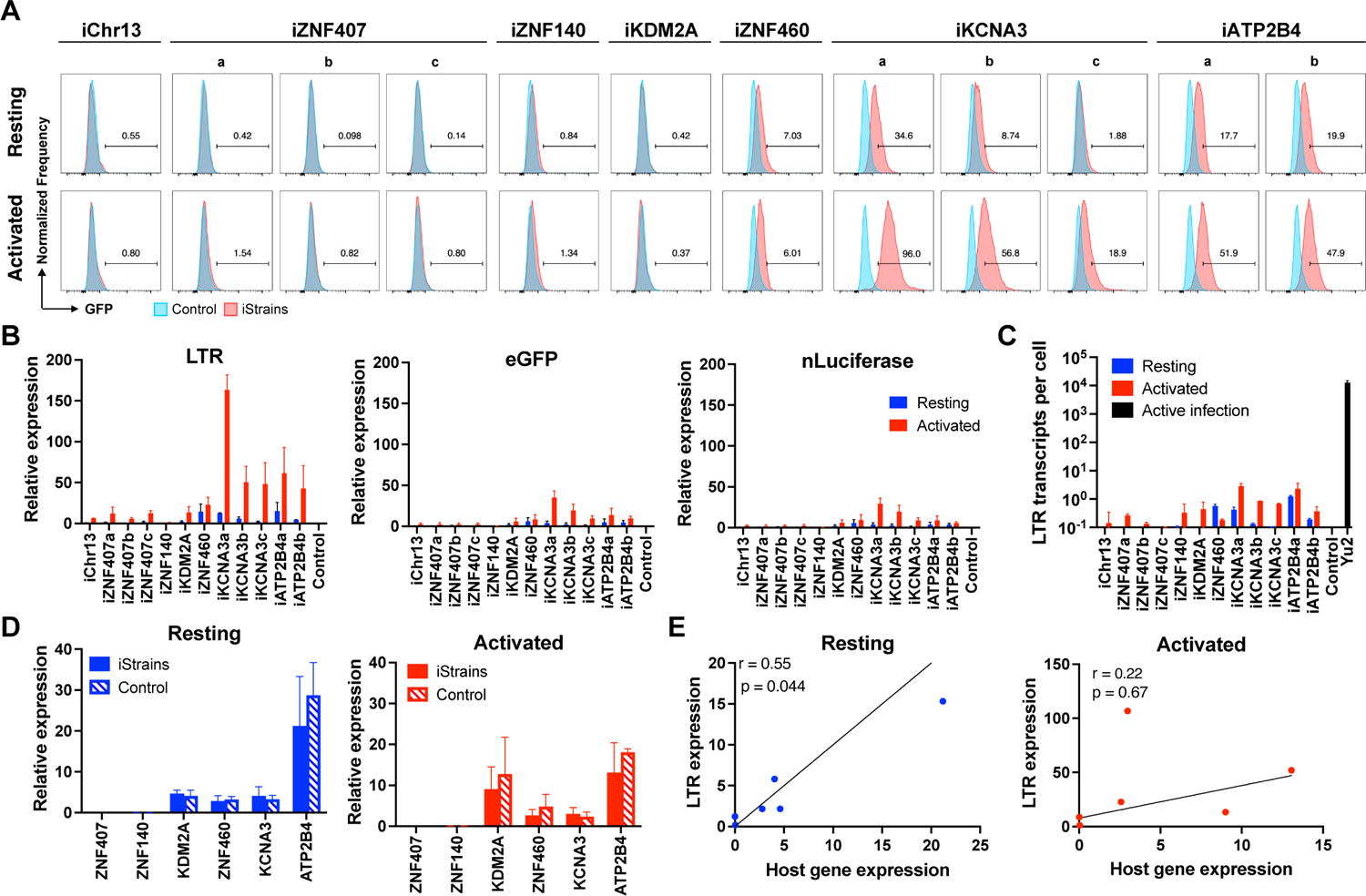
Jurkat reporter lines. **(A)** Histograms show GFP fluorescence (x-axis) per normalized counts (y-axis) for each integration site studied (iStrains, red) and control (blue), in both resting (upper panel) and PMA/ionomycin-activated (lower panel) conditions. **(B)** Graphs show relative LTR (left panel), eGFP (middle panel) and nLuciferase (right panel) expression assessed by qPCR under resting (blue) and PMA/ionomycin-activated (red) conditions. Bars represent the mean relative expression from two independent assays (biological replicates) ± standard deviation. **(C)** LTR transcripts per cell (y-axis), determined by qPCR, for each integration-positive clone and control, under resting (blue) and activated (red) conditions. CD4^+^ T cells infected with HIV-1_Yu2_ served as positive control (black bar). Bars represent the mean of two independent assays (biological replicates) ± standard deviation. **(D)** Relative expression determined by qPCR for host genes neighboring their respective reporter proviruses (iStrains, full bars) and control (stripped bars), under resting (left panel, blue) and PMA/ionomycin-activated (right panel, red) conditions. Bars represent the mean relative expression of two independent assays (biological replicates) ± standard deviation. Expression of host gene was averaged across multiple clones to the same reporter integration site. **(E)** Correlation between the relative expression of LTR (y-axis) for each Jurkat clone and their respective host gene (x-axis), averaged across multiple clones to the same reporter integration site, under resting (left panel, blue) and PMA/ionomycin-activated (right panel, red) conditions. Person’s correlation coefficients, r, and two-tailed p values were computed for each condition.

**Figure 3.**
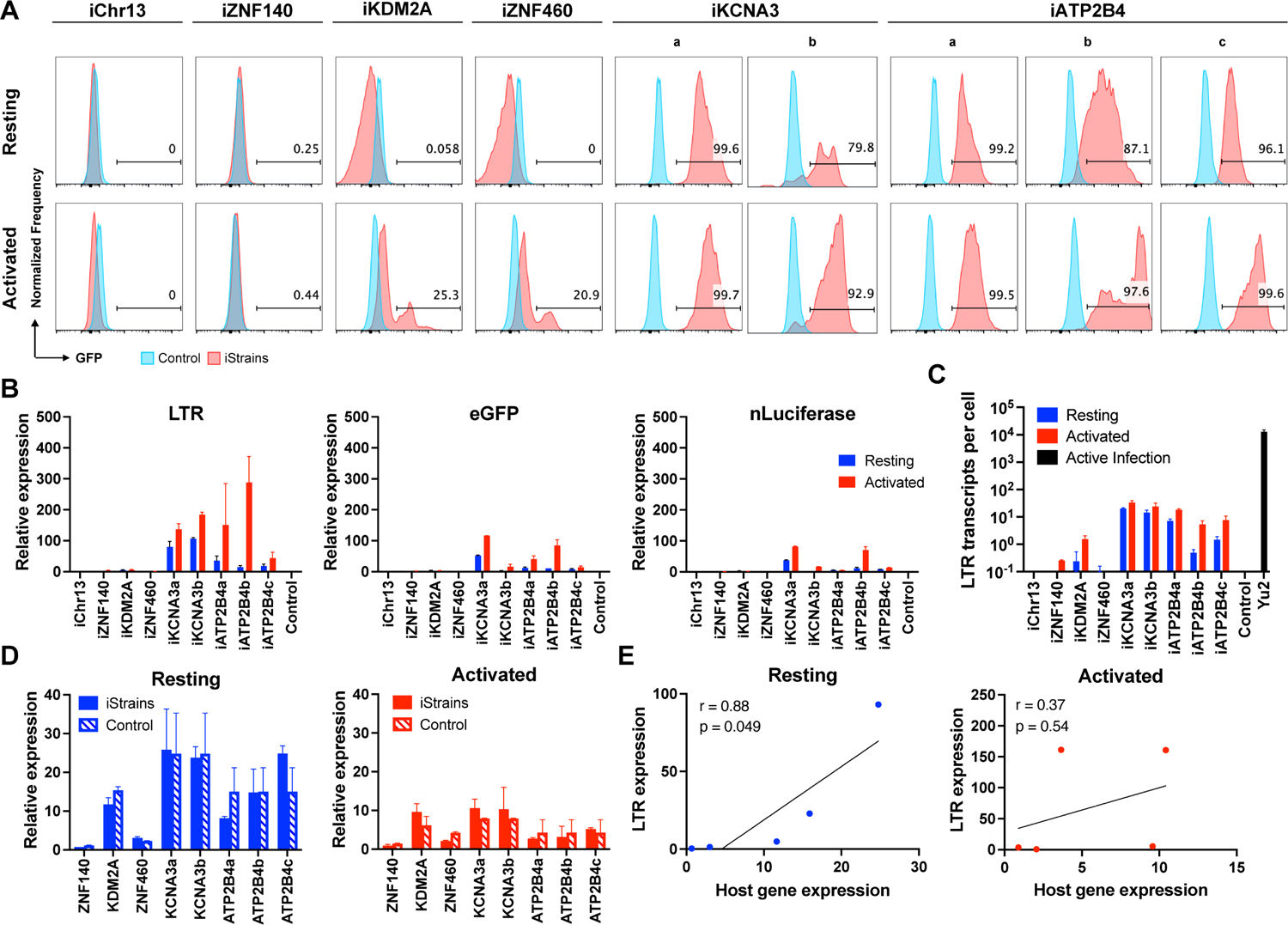
Primary CD4^+^ T cell reporter lines. **(A)** Histograms show GFP fluorescence (x-axis) per normalized counts (y-axis) for each integration site studied (iStrains, red) and control (blue), in both resting (upper panel) and CD3/CD28-activated (lower panel) conditions. **(B)** Graphs show relative LTR (left panel), eGFP (middle panel) and nLuciferase (right panel) expression assessed by qPCR under resting (blue) and CD3/CD28-activated (red) conditions. Bars represent the mean relative expression from two independent assays (biological replicates) ± standard deviation. **(C)** LTR transcripts per cell (y-axis), determined by qPCR, for each integration-positive clone and control, under resting (blue) and CD3/CD28-activated (red) conditions. CD4^+^ T cells infected with HIV-1_Yu2_ served as positive control (black bar). Bars represent the mean of two independent assays (biological replicates) ± standard deviation. **(D)** Relative expression determined by qPCR for host genes neighboring their respective reporter proviruses (iStrains, full bars) and control (stripped bars), under resting (left panel, blue) and CD3/CD28-activated (right panel, red) conditions. Bars represent the mean relative expression of two independent assays (biological replicates) ± standard deviation. **(E)** Correlation between the relative expression of LTR (y-axis) for each primary T cell clone and their respective host gene (x-axis), averaged across multiple clones to the same reporter integration site, under resting (left panel, blue) and activated (right panel, red) conditions. Person’s correlation coefficients, r, and two-tailed p values were computed for each condition.

### Proviral RNA expression

Transcription of intact HIV-1 proviruses in the latent reservoir has been documented by sensitive single copy PCR assays (Einkauf et al., 2022). However, the assay involves multiple rounds of polymerase amplification and is not quantitative. Thus, the amount of transcription at the sites of intact latent proviral integration, how it compares to productive infection, and how it relates to the site of integration has not been examined.

We performed quantitative PCR (qPCR) assays to determine the level of transcription of integrated proviral reporters under resting and activated conditions in Jurkat and primary CD4^+^ T cell lines (Fig. 2 B and 3 B and Fig S1 C). Under resting conditions, reporter LTR expression was either very low at Chr13, *ZNF407*, *ZNF140*, *KDM2A* and *ZNF460* sites in Jurkat cell lines (Fig. 2B) or undetectable in primary CD4^+^ T cells (Fig. 3B). eGFP and nLuciferase were transcribed at lower levels than the LTR and only detectable in reporters integrated at *KCNA3* and *ATP2B4* in primary CD4^+^ T cells suggesting incomplete transcript elongation (Fig. 2 B and 3 B). Activation with PMA/Ionomycin or anti-CD3/28 antibodies increased LTR mRNA expression to measurable levels at all sites of integration except *ZNF140* in Jurkat cells (Fig. 2 B and Fig S1 C). In contrast, only reporters integrated into *KCNA3* and *ATP2B4* showed increased LTR expression in primary CD4^+^ T cells (Fig. 3 B). However, even after activation the level of reporter LTR expression, quantified as the number of transcripts per cell, was 10^3^-10^4^ and 10^2^-10^3^-fold lower than HIV-1_YU2_ infected CD4^+^ T cells in Jurkat and primary CD4^+^ T cells respectively (Fig. 2 C and 3 C). Thus, the results in Jurkat and primary CD4^+^ T cells are generally congruent, but reporter expression levels are lower in Jurkat cell lines.

### Host gene expression and chromatin architecture

To determine how proviruses affect nearby host genes we measured their expression by qPCR under resting and activated conditions. In all cases in Jurkat and primary CD4^+^ T cells, host gene mRNA expression was not significantly altered by reporter integration (Fig. 2 D and 3 D). Similar results were obtained using a different reporter that included 2 LTRs (Fig. S2). Under resting conditions, reporter LTR expression in Jurkat and primary CD4^+^ T cells was directly but not precisely proportional to the level of expression of the host gene (Fig. 2 E and 3 E). For example, in *ZNF140* and *ZNF460* reporter LTRs are expressed at very low levels in both Jurkat and primary CD4^+^ T cells. However, *KDM2A* is expressed at relatively high levels in primary CD4^+^ T cells but the reporter is silent (Fig. 3 E). Notably, there was no significant correlation between integration site and proviral transcription after activation suggesting that under these conditions regulation is no longer dominated by the host gene.

To determine how proviral integration might alter genome accessibility in or near the site of integration we performed an assay for transposase-accessible chromatin with sequencing (ATAC-seq) on all Jurkat and primary CD4^+^ T cell lines. The reporter LTR was accessible in all cell lines except in *ZNF140* and, surprisingly, *ZNF460* in Jurkat cells and the non-genic region in Chr13 in primary CD4^+^ T cells (Fig. 4 A and B). Thus, even in the absence of measurable transcription the reporter remained accessible in most sites of latent integration. Notably, except for *KCNA3* in primary CD4^+^ T cells, the presence of the reporter did not alter neighboring genome chromatin accessibility irrespective of proviral transcription under resting or activated conditions (Fig. 4 C and Fig. S3 and S4). We conclude that HIV-1 LTR reporter integration at authentic latent sites in Jurkat or primary CD4^+^ T cells does not alter local gene expression and only occasionally impacts local chromatin accessibility.

**Figure 4.**
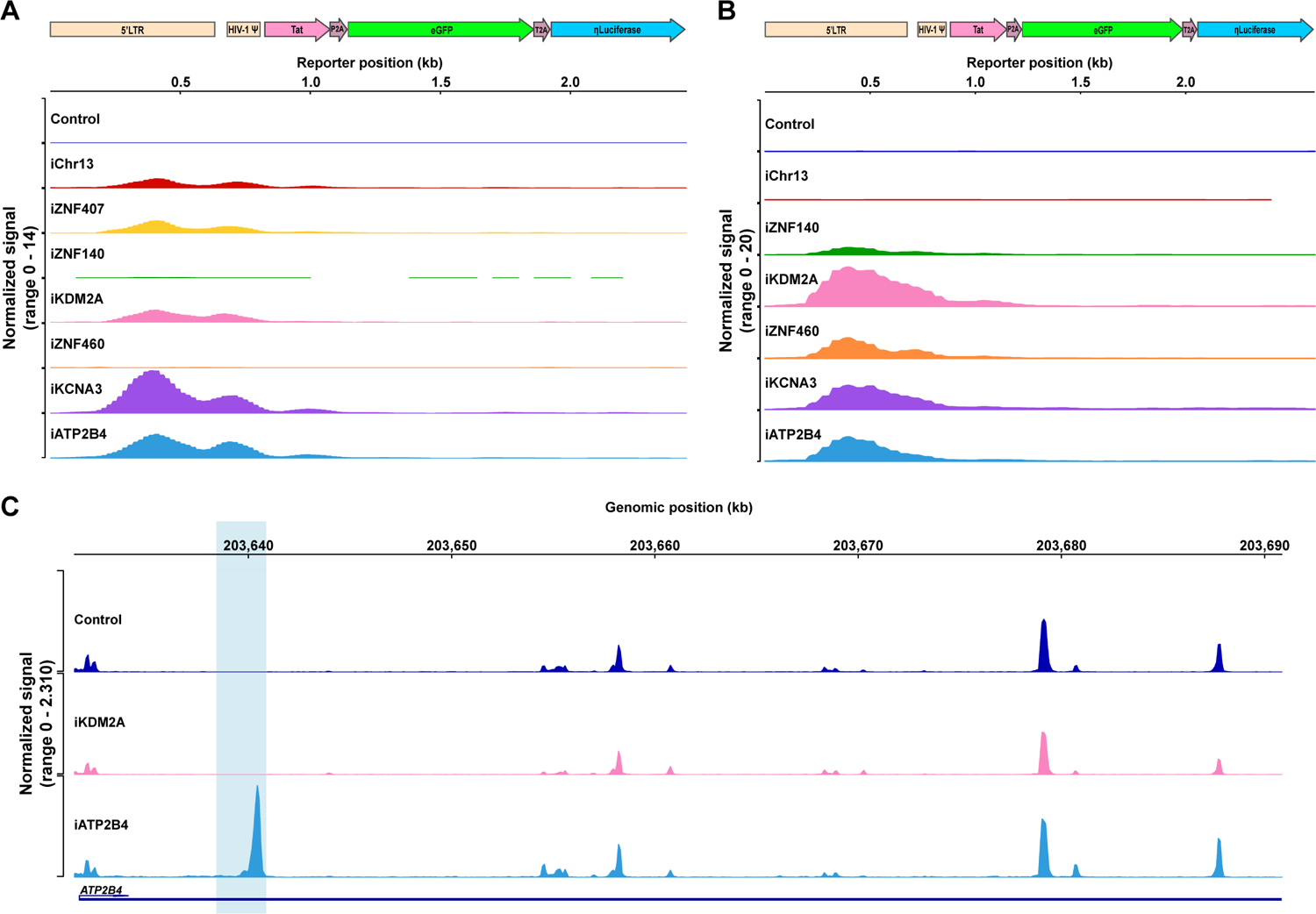
Chromatin accessibility. **(A-B)** Chromatin accessibility in Jurkat **(A)** and primary CD4^+^ T cell clones **(B)** as measured by ATAC-seq for the reporter construct at each integration site and control under resting conditions **(C)** Graph shows ATAC-seq for *ATP2B4* encompassing the reporter integration site in control, and clones that carry the reporter in *KDM2A*, *ATP2B4*. Blue shading indicates the site of reporter integration. Graphs were generated by averaging the normalized reads from three technical replicates for each clone.

### Intact latent HIV-1 proviruses

CD4^+^ T cells carrying intact latent HIV-1 proviruses are rare and express no singular cell surface marker that distinguishes them from T cells other than their unique T cell receptor (TCR) (Cohn et al., 2018; Collora et al., 2022; Einkauf et al., 2022; Weymar et al., 2022; Sun et al., 2023; Wu et al., 2023). To examine HIV-1 expression in authentic latent reservoir cells we enriched CD4^+^ T cells carrying replication competent latent proviruses by means of their specific TCRs (Weymar et al., 2022) (#603 and #5104 with HIV-1 integrated into *ZNF486* and *ATP2B4*). The sorted cells were then expanded under limiting dilution conditions in the presence of irradiated feeder cells and anti-retroviral drugs (ARVs). The presence and frequencies of the infected clones in the expanded cells lines was verified by PCR assays for their specific TCRs and HIV-1 sequences (Fig. 5 A). Cell lines were obtained in which at least half of the cells represented the T cell clones of interest with intact proviruses integrated into *ZNF486* and *ATP2B4* genes.

**Figure 5.**
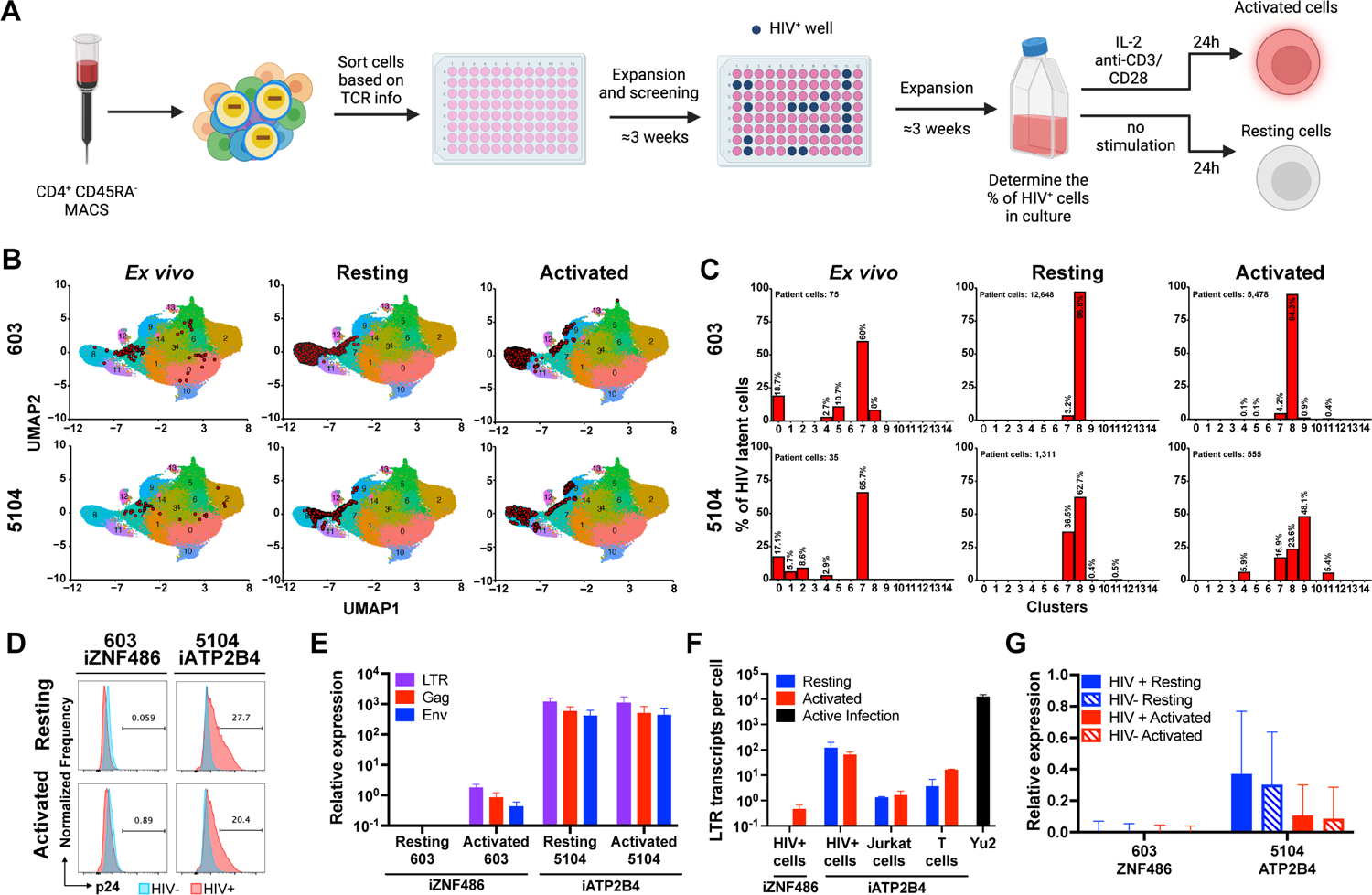
HIV-infected cells from people living with HIV. **(A)** Schematic representation of the methods used to grow out HIV-1 infected cells from ART suppressed individuals (Weymar et al., 2022). Created with BioRender.com. **(B-C)** Uniform manifold approximation and projection (UMAP) of 10X single cell gene-expression data showing the position of the cells expressing the latent clones’ specific TCR (red dots) **(B)** and the fraction of latent cells in each cluster **(C)**, for participants 603 (upper panels) and 5104 (lower panels) from *ex vivo* cells (left panels, data from Weymar et al., 2022), and cultured cells under resting (middle panels) and activated (right panels) conditions. **(D)** Histograms show HIV-1 Gag p24 expression in HIV+ cells (red) and non-infected cells (blue) of cultures derived from 603 and 5104, under resting (upper panel) and activated (lower panel) conditions. **(E)** Relative expression of LTR (purple bars), Gag (red bars) and Env (blue bars) by qPCR for 603 and 5104 HIV+ cells under resting and activated conditions. Bars represent the mean relative expression of two independent assays (biological replicates) ± standard deviation. **(F)** LTR transcripts per cell determined by qPCR, in cells from 603 and 5104 and in Jurkat and primary T cells reporter lines with proviruses integrated into iATP2B4, under resting (blue) and activated (red) conditions and HIV-1_YU2_ controls. Bars represent the mean of two independent experiments (biological replicates) ± standard deviation. **(G)** Relative expression determined by 10X Genomics single cell mRNA sequencing of host genes neighboring HIV-1 proviral integration (*ZNF486*, participant 603 and *ATP2B4,* participant 5104) in HIV-infected (HIV+, full bars) and non-infected (HIV-, stripped bars) cells from the same cell population, under resting (blue bars) and activated (red bars) conditions. Bars represent the mean of the respective population from one assay ± standard deviation representing population variance.

Primary CD4^+^ T cells carrying intact latent proviruses isolated directly from individuals 5104 and 603 are enriched in T cell populations with specific transcriptional profiles (Weymar et al., 2022). To determine the transcriptional profile of cultured lines of CD4^+^ T cells carrying integrated HIV-1 proviruses in *ZNF486* and *ATP2B4* genes we performed single cell mRNA sequencing experiments using the 10X Genomics platform. Specific TCR expression was used to identify the latent cells and map them onto our previous data set using a cut-off of 95% confidence (Fig. 5 B and C, and materials and methods) (Weymar et al., 2022). Cultured and *ex vivo* cells containing latent HIV-1 proviruses showed closely related transcriptional profiles (Fig. 5 B and C).

The enriched populations of resting (3 weeks after activation) and activated (24 hours after anti-CD3 and -CD28 monoclonal antibody stimulation) CD4^+^ T cells carrying the 5104 and 603 HIV-1 proviruses were initially examined for p24 expression by flow cytometry (Fig. 5 D). Like the reporters in Jurkat and CD4^+^ cell lines, the HIV-1 provirus integrated into *ATP2B4*, an actively transcribed gene, showed expression under both resting and activated conditions (Fig. 5 D). In contrast, the provirus integrated into *ZNF486*, a gene expressed at only low levels, failed to show detectable p24 expression under resting or activated conditions (Fig. 5 D).

HIV-1 LTR, Gag and Env mRNA expression were measured by qPCR corrected for the fraction of infected cells in the culture (Fig. 5 E, and materials and methods). The HIV-1 provirus integrated into *ATP2B4* showed similar levels of expression of LTR, Gag and Env under resting and activated conditions. In addition, we found multiple spliced transcripts associated with productive infection in single cell transcriptome analysis (Fig. S5). In contrast, the HIV-1 provirus integrated into *ZNF486*, only showed LTR, Gag and Env transcripts after activation (Fig. 5 E). When compared to productive infection with HIV-1_YU2_, the proviruses in *ATP2B4* and *ZNF486* were expressed at >100 and >27,000-fold lower levels respectively (Fig. 5 F). When compared to reporter proviruses integrated into *ATP2B4* the level of LTR expression obtained from cultured CD4^+^ T cells carrying latent HIV-1 was 10 and 100-fold higher than in primary CD4^+^ T cells and Jurkat cell respectively. We conclude that authentic latent HIV-1 proviruses integrated at different sites in the genome of CD4^+^ T cells show different levels of expression and ability to respond to activation. In addition, proviruses integrated at latent sites are expressed at far lower levels than in productive infection.

To determine whether the latent HIV-1 proviruses in the cultured cells altered neighboring host gene transcription (*ATP2B4* and *ZNF486*) their expression was compared with non-infected cells from the same culture in the 10X Genomics data set. We found no significant difference in expression of genes neighboring the integrated HIV-1 provirus between infected and non-infected cells irrespective of activation (Fig. 5 G).

## Discussion

HIV-1 infection typically results in the death of the infected cell. However, CD4^+^ T cells harboring intact chromosomally integrated HIV-1 proviruses can persist for years in individuals that are virally suppressed on ART (Dufour et al., 2021; Lichterfeld et al., 2022; Siliciano and Siliciano, 2022). The provirus is silent in some of these latent cells, which may explain why they persist, but intact proviral transcription has been detected using sensitive but non-quantitative PCR methods (Einkauf et al., 2022). We have examined HIV-1 proviral transcription at sites of authentic intact latent proviral integration in Jurkat cell lines, primary CD4^+^ T cells, and CD4^+^ T cell lines derived from PLWH. The data reveal that in resting, but not activated conditions, the transcriptional activity at the site of intact proviral integration is correlated to the level of proviral gene expression. Notably the intact proviruses studied had little or no detectable impact on neighboring gene expression.

The reservoir of CD4^+^ T cells that harbor latent proviruses is both heterogeneous and dynamic. It is composed of clones of CD4^+^ T cells that expand and contract in a manner that is in part dependent on antigenic stimulation and trophic cytokines required for CD4^+^ T cell longevity (Demoustier et al., 2002; Douek et al., 2002; Chomont et al., 2009; Saleh et al., 2011; Jones et al., 2012; Cohn et al., 2015; Lorenzi et al., 2016; Bui et al., 2017; Hosmane et al., 2017; Lee et al., 2017; Pinzone et al., 2019; Scheerder et al., 2019; Antar et al., 2020; Mendoza et al., 2020; Simonetti et al., 2021; Cho et al., 2022). Over time the measurable reservoir contracts with an initial half-life of approximately 4 years and becomes more clonal (Siliciano et al., 2003; Crooks et al., 2015; Bachmann et al., 2019; Peluso et al., 2020; Cho et al., 2022). Larger clones of CD4^+^ T cells harboring intact proviruses appear to be more difficult to reactivate (Lorenzi et al., 2016) and their pro-viruses are preferentially found in transcriptionally inactive parts of the genome (Einkauf et al., 2019; Huang et al., 2021). Sensitive PCR-based methods indicate that this shift in the reservoir is associated with retroviral transcription with preferential survival of silent clones (Einkauf et al., 2022). Thus, the transcriptional status of the intact provirus appears to be an essential element in determining its longevity. Our experiments indicate that there is a great deal of variability in the level of latent proviral expression that tracks with integration site and cell type. How the site of integration of intact latent proviruses or its transcription might alter latent cell longevity is not known, but the mRNA or protein products of the virus could have a direct effect on the cell or indirectly engage host immunity.

We found little effect on transcription of neighboring genes by reporter proviruses or HIV-1 integrated at 8 sites of intact latent proviral integration in Jurkat cell lines, primary CD4^+^ T cells or CD4^+^ T cells obtained from PLWH and grown in culture. Initially, intact latent proviruses are preferentially found in introns of genic regions, in the opposite transcriptional orientation to the transcriptional start site (Einkauf et al., 2019; Huang et al., 2021). Over time in chronically infected individuals under ART and elite controllers, intact latent proviruses accumulate in ZNF genes and non-genic regions (Einkauf et al., 2019; Jiang et al., 2020; Huang et al., 2021). Our collection of integration sites mirrors this selection and therefore the observation that intact latent proviruses have little measurable effect on the neighboring genome may be biased by the origins of the sample set. Nevertheless, the integrations examined include unique proviruses found only once and others found in expanded clones.

In contrast to proviral influence on neighboring genes, the transcriptional status of the genes at integration sites were directly correlated to LTR transcription. Notably, the correlation between the proximal host gene expression and LTR transcription was lost upon T cell activation. In some instances, proviral transcription was increased even as proximal host gene expression was downregulated. These results are consistent with a Tat-independent phase during which basal expression depends mostly on host factors and a second, Tat-dependent positive feedback loop that promotes proviral transcription upon activation (Jordan et al., 2001). However, proviruses integrated in highly repressed genomic regions, i.e. *ZNF140* or in non-genic regions, remained silent after T cell stimulation. Our findings are consistent with observations made with randomly integrated reporter proviruses in cells lines that the level of HIV expression depends on the DNA integration site (Jordan et al., 2001; Jordan, 2003; Lewinski et al., 2005; Skupsky et al., 2010). However, our sample size is limited, and larger and more diverse group of authentic latent integration sites is necessary to understand the influence of integration site on proviral expression precisely. Moreover, our reporters are missing components of the provirus that can impact the host cell and promote proviral survival and transcription (Malim and Emerman, 2008; Cesana et al., 2017; Pinzone et al., 2019; Yeh et al., 2020). Their absence may in part explain why we observe no effects on host transcription or why transcription from the indicator lines is lower than the authentic latently infected cell lines carrying an intact provirus in the same genomic location.

Notably, we found somewhat variable levels of expression between clones carrying the same integration. In addition, only a fraction of CD4^+^ T cells expressed the reporters integrated in *KDM2A* and *ZNF460* after activation, Our observations are consistent with the finding that HIV-1 transcription is inherently stochastic (Ne et al., 2018; Mori and Valente, 2020) which may result in varying levels of gene expression in single cells within a clonal population. This effect may be further compounded by the stochasticity of host gene transcription in the clones (Raj and Oudenaarden, 2008). This variability may in part explain why activation of proviruses *in vitro* requires multiple rounds of stimulation (Hosmane et al., 2017).

We studied 2 cell lines derived directly from latently infected cells obtained from people living with HIV-1. Proviral transcription was detected in cells where HIV-1 was integrated in a highly expressed gene, *ATP2B4*, but not in cells carrying a provirus integrated in *ZNF486*, a gene expressed at low levels. Although limited to 2 examples the results are congruent with our observations using indicator proviruses in Jurkat and primary CD4^+^ T cells. In all cases the levels of HIV-1 transcription from sites of intact latent integration were at least 2 orders of magnitude below the levels found in productively infected cells. Although insufficient to kill all the infected cells, the low level of proviral transcription from the *ATP2B4* integration was sufficient to produce infectious virus (data not shown). Thus, latent CD4^+^ T cells carrying an HIV-1 provirus integrated into *ATP2B4* are also likely to produce enough viral protein to be visible to the immune system.

In conclusion, the direct relationship between integration site and transcriptional regulation provides a road map for understanding how individual proviruses in the reservoir are selected against and how they contribute to viral rebound.

## Materials and methods

### Participant material and cell lines

Jurkat cells were procured from ATCC (clone E6-1, TIB-152). Primary CD4^+^ T cells were isolated from frozen PBMCs of a healthy individual, while latent HIV+ cells were isolated as described below from frozen PBMCs of participants of a previous study. Participant cohort information is described in Table S1 and in a previous publication (Weymar et al., 2022).

### CRISPR RNA (crRNA) design

crRNAs were designed using the Integrated DNA Technologies crRNA design tool (http://www.idtdna.com) and synthesized by Integrated DNA Technologies as Alt-R CRISPR/Cas9 RNAs. crRNA sequences are listed in Table S2.

### Homology-Directed Repair Template (HDRT) preparation and AAV production

HDRT sequences (Table S1), were cloned into a pAAV vector using Gibson assembly technique (Gibson et al., 2009) with the NEBuilder^®^ HiFi DNA Assembly Master Mix (New England Biolabs, cat. E2621L). AAVs carrying each HDRT were produced as previously described (Hartweger et al., 2023). Our strategy is optimized for targeting efficiency and would not accommodate a full-length provirus that cannot be incorporated into AAV.

### Nucleofection of Jurkat and primary human CD4^+^ T cells

Genome editing was achieved through a previously described combination of CRISPR targeting and AAV-mediated homologous recombination (Martin et al., 2019).

Primary CD4^+^ T cells were enriched from healthy donor cryopreserved PBMCs by negative selection (CD4^+^ T Cell Isolation Kit human, Miltenyi Biotec, cat. 130-096-533) and activated for 72h with 10 µg/mL of plate-bound Ultra-LEAF™ purified anti-human CD3 monoclonal antibody (clone OKT3, BioLegend, cat. 317325) and 5 µg/mL of soluble Ultra-LEAF™ purified anti-human CD28 monoclonal antibody (clone 28.2, BioLegend, cat. 302933) in R10 medium [RPMI-1640 medium supplemented with 10% heat-inactivated FBS, 10 mM Hepes, antibiotic-antimycotic (1x) and 2 mM L-glutamine (all from Gibco)] supplemented with recombinant human IL-2 (50 U/mL; Roche, cat. 10799068001).

For a 100 µL transfection, 1 µL of 200 µM of gene-specific crRNA and 1 µl 200 µM trans-activating CRISPR RNA (tracrRNA) in duplex buffer (all from Integrated DNA Technologies) were combined, denatured at 95°C for 5 min, and renatured for 5 min at room temperature. Then, 5.6 µl PBS and 2.4 µl 61 µM Cas9 (Alt-R™ *S.p.* Cas9 Nuclease V3, Integrated DNA Technologies, cat. 1081058) were added and the mixture incubated for 20 min. Finally, 4 µl 100 µM electroporation enhancer in duplex buffer was added to 10 µl of the RNPs and the mixture incubated for a further 1–2 min.

Nucleofection of Jurkat cells was performed using SE Cell Line 4D-Nucleofector™ X Kit L (Lonza, cat. V4XC-1012), while the P3 Primary Cell 4D-Nucleofector™ X Kit L (Lonza, cat. V4XP-3024) was used for primary T CD4^+^ T cells, as previously described (Hultquist et al., 2019). After nucleofection cells were transferred into a 12-well plate containing R10 medium and the corresponding HDRT-carrying AAV.

### Jurkat and primary human CD4^+^ T cells

Three days post nucleofection, cells were stained with Fixable Viability Dye eFluor 780 (Invitrogen, cat. 65-0865-14) and single cell sorted using a FACSymphony S6 using FACSDiva software (version 9.5.1, BD Biosciences) into R10 (Jurkats) or activation medium (R10 + 50 U/mL IL-2, 0.1 µg/mL of anti-CD3 and -CD28 monoclonal antibodies) plus feeder cells (1×10^6^/mL NK and CD8+ T cell-depleted, 5000 rad irradiated allogeneic PBMCs). Three weeks after sorting, a small fraction of the cells was collected, genomic DNA extracted (QuickExtract™ DNA Extraction Solution, Lucigen, cat. QE09050) and the remainder restimulated with fresh medium and cultured for an additional 14 days (Jones et al., 2016).

Clones were screened for integration with combinations of HDRT- and gene-specific primers, covering the entire construct and the integration sites (Table S2). PCRs were performed using Phusion Green Hot Start II High-Fidelity DNA Polymerase (ThermoFisher Scientific, cat. F537S) and Sanger sequencing of gel extracted amplicons performed (Azenta Life Sciences).

### CD4^+^ T cells from individuals on ART

CD4^+^ T cells harboring intact latent HIV-1 proviruses were enriched using their T cell receptors (Weymar et al., 2022). CD4^+^ T cells from participants 603 (Rockefeller University IRB-approved Protocols MCA-0866/TSC-0910) and 5104 (Protocols MCA-0965/TSC-0910) were isolated by magnetic separation using CD4^+^ T Cell Isolation Kit, human (Miltenyi Biotec, cat. 130-091-155) and CD45RA MicroBeads, human (Miltenyi Biotec, cat. 130-045-901) and prepared for cell sorting exactly as described (Weymar et al., 2022). Cells from 603 (CD4^+^TRBC1^+^TRBV19^+^) and 5104 (CD3^+^CD4^+^TRBC1^+^TRBV) were sorted into 96 well “U” bottom plates (5 cells per well) containing 200 µL of activation media (composition described above), plus feeder cells, supplemented with the following ARVs: 1µM Tenofovir, 1µM Emtricitabine, 1nM Nevirapine and 10 µM T-20. Cell sorting was performed on FACSymphony S6 using FACSDiva software (version 9.5.1, BD Biosciences) and data was analyzed using FlowJo (version 10.10.0, BD Biosciences). The culture media was replaced twice a week to replenish ARVs and avoid new infections.

Three weeks after sorting, cells were screened for the presence of the HIV clone of interest by Env PCR and sequencing (Salazar-Gonzalez et al., 2008). Cell lines containing clones of interest were expanded in a T25 culture flask in 10 mL of activation medium. The frequency of latent cells in the cultures was determined three weeks after expansion by sorting single cells into 10 µL of RLT buffer (Qiagen, cat 79216) using Agencourt RNAClean XP and magnetic beads (Beckman Coulter, cat. A63987) for DNA preparation and Env PCR.

### Activation of cultured cells

Jurkat cells were activated in R10 medium supplemented with PMA (25 ng/mL, Sigma-Aldrich, cat. P1585) and ionomycin (1 μg/mL, Sigma-Aldrich, cat. I9657) or R10 supplemented with anti-human CD3 and CD28 antibodies (as above) for 24 h.

Primary CD4^+^ T cells carrying reporter constructs or those obtained from individuals 603 and 5104 or controls, 3-4 weeks after the last stimulation in activation medium plus feeder cells, were stimulated with activation medium (described above) for either 24 hours or one week. In parallel, cells were maintained in resting conditions by culturing in simple R10 or R10 supplemented with 10 ng/mL of recombinant human interleukin-7 (IL-7, R&D Systems, cat. BT-007-010) for one week.

### Flow cytometric analysis

T cell clones were assessed for GFP expression by flow cytometry. Cells were stained with Fixable Viability Dye eFluor 780 and the following monoclonal antibodies: TexasRed-PE conjugated anti-human CD3 (clone 7D6, Invitrogen, cat. MHCD0317); PerCP/Cyanine5.5 conjugated anti-human CD4 (clone OKT4, BioLegend, cat. 317428); BrillianViolet conjugated anti-human CD25 (clone BC56, BioLegend, cat. 302634).

HIV Gag p24 expression was evaluated by flow cytometry in CD4^+^ T cells derived from participants 603 and 5104. Cells were fixed and permeabilized using BD Cytofix/Cytoperm™ Fixation/Permeabilization Kit (BD Biosciences, cat. 554714) and stained for Gag p24 protein with anti-HIV-1 core antigen monoclonal antibody (clone KC57, Beckman Coulter, cat. 6604665).

Cells were analyzed in a FACS Symphony A5 flow cytometer running FACS Diva software (version 8.5, BD Biosciences) and data was analyzed using FlowJo (version 10.10.0, BD Biosciences).

Only live cells, CD25^high^ (activation conditions) or CD25^low^ (resting conditions) were considered for measuring GFP or p24 fluorescence.

### qPCR

RNA was purified from the bulk clonal populations using the RNeasy® Plus Micro kit (Qiagen, cat. 74034) and converted to cDNA using SuperScript^TM^ III Reverse Transcriptase (Invitrogen, cat. 18080-093), using a combination of random primers (Invitrogen, cat. 48190011) and LTR-specific primers for reporter primary cells, plus primers to polyA, nef, tat-rev and pol for HIV+ cells (Table S1) (Einkauf et al., 2022). qPCR was performed on cDNA using TaqMan™ Fast Advanced Master Mix for qPCR (Applied Biosystems, cat. 4444558) with 500 nM primers and 250 nM probe plus 1 µL of cDNA. For relative quantification of expression, normalization was performed against the *PPIA* gene. Primers and probes used for quantification of LTR, eGFP and nanoLuciferase were from Integrated DNA Technologies (Table S1). For host gene expression we used pre-designed PrimeTime qPCR Assays (Integrated DNA Technologies): *ZNF407* – Hs.PT.58.4027445, *ZNF140* – Hs.PT.58.39945645, *KDM2A* – Hs.PT.58.815648, *ZNF460* – Hs.PT.58.24604991, *KCNA3* – Hs.PT.58.2591067.g, *ATP2B4* – Hs.PT.56a.26489531.g and *PPIA* - Hs.PT.58v.38887593.g, PIKFYVE - Hs.PT.58.26959495 and Hs.PT.58.40848161, FIZ1 - Hs.PT.58.2780215 and Hs.PT.58.5062508. HIV specific primers for poly-A, nef, tat-rev, pol, and long LTR were as previously described (Palmer et al., 2003; Gaebler et al., 2019; Einkauf et al., 2022) and Table S**2**. Relative expression was calculated as 2^-(ct*target*-ct*ref*)^. Number of LTR transcripts were enumerated in a qPCR reaction, using the same LTR primers as above. A standard curve of 10-fold serial dilutions a plasmid carrying a LTR sequence was created by plotting known DNA copy number in each dilution to their respective Ct value. Ct values measured for LTR in each sample were inputted in the standard curve and the transcript copy number thus obtained was divided by the number of cells, or infected cells for HIV+ cell cultures, equivalent to input volume of cDNA in the reaction to obtain the number of LTR transcripts per cell.

### ATAC-seq

Dead cells were removed from culture using the Dead Cell Removal Kit (Miltenyi Biotec, cat. 130-090-101). Then, 100,000 Jurkat cells or primary CD4^+^ T cells were used per reaction. For primary T cells, each clone was sampled in both resting and activated conditions as described above. Three replicates were performed per condition. Cells were lysed and DNA tagmented, purified and amplified using an ATAC-Seq kit (Active Motif, cat. 53150). The libraries were sequenced using Illumina NextSeq 550 paired-end sequencing.

The human genome assembly hg38 was modified adding HIV inserts at specified loci using custom scripts. Subsequently, all libraries were mapped to this modified genome using bowtie2 with the parameters --local --very-sensitive-local. Duplicated reads were then removed with Picard MarkDuplicates with default parameters. Regions with enriched signal throughout the genome were obtained using macs2 for peak calling with the parameters --format BAMPE --nomodel –nolambda --cutoff-analysis, genome blacklist regions (https://github.com/Boyle-Lab/Blacklist) were removed (Amemiya et al., 2019). Differentially accessible regions were identified using edgeR.

### 10X Genomics

10X Genomics gene expression and V(D)J libraries were generated with the Chromium Single Cell 5′ Library & Gel Bead Kit (10X Genomics, cat. PN-1000014) and Chromium Single Cell V(D)J Enrichment Kit, Human T Cell (10X Genomics, cat. PN-1000005) as described in the 10X Genomics protocol. The 5’ expression library and the V(D)J library were sequenced with NovaSeq 6000 S1 (100 cycles) (Illumina, cat. 20012865).

### Single-cell RNA-seq (scRNA-seq) and single-cell TCR-seq processing

Single-cell RNA-seq binary base call (BCL) files underwent demultiplexing and were transformed into FASTQ files using BCLtoFastq, followed by alignment against a modified hg38 that includes specific HIV sequences for each subject, utilizing CellRanger (v7.2.0). Analysis was performed in R studio using Seurat (v5.0.1). Cells exhibiting mitochondrial content over 10% and/or feature counts outside the 200 to 2,500 range were excluded. Sample batches were merged, then normalized and scaled using SCTransform. Single-cell TCR-seq FASTQ files were aligned to the standard CellRanger VDJ reference using CellRanger (v7.2.0). The resulting contig annotations were filtered and examined in R studio. Cells harboring TCRs identical to those found by Weymar and colleagues (Weymar et al., 2022), were classified as latent cells, whereas those with diverse TCRs were identified as non-infected cells.

### Mapping scRNA-seq from cultured cell to previously published UMAP

The UMAP from a previous report (Weymar et al., 2022) served as the reference, and the cultured cells from each subject were anchored and mapped using the FindTransferAnchors and MapQuery functions from Seurat (reference.reduction = “pca”, dims = 1:30, reduction.model = umap, refdata = clusters). Only cells with a prediction score of 0.95 or higher for label transferring were selected to create Fig 5 B.

### Statistical analysis

Person’s correlation coefficients, r, and two-tailed p values were computed for the relationship between the host gene and reporter LTR expression. Statistical analyses were performed in GraphPad PRISM version 10. Analysis of ATAC-Seq and single cell sequencing data were processed and analyzed using R studio version 2023.12.1+402 running R version 3.3.0.

### Online supplementary material

Table S1 details the clinical characteristics of the participants of previous studies whose cells were used in the current work. Table S2 lists crRNA sequences, annotated HDRT sequences and primer sequences for the various assays.

## Acknowledgments

We thank all study participants who devoted time to our research, the Rockefeller University Hospital Research support office and nursing staff, all members of the Nussenzweig laboratory for discussions, and M. Jankovic, G. Scrivanti and D. Solari for laboratory support as well as L. Cohn for discussions and for critically reading the manuscript. We thank C. Zhao, H. Duan, C. Lai, and S. Huang from the Genomics Resource Center at the Rockefeller University for preparing and sequencing the 10x Genomics and ATAC-seq libraries. We also thank J.P. Truman and K.M. Gordon for operating the cell sorters. This work was supported by the National Institutes of Health (grants UM1 AI100663 and R01AI129795 to M.C.Nussenzweig), REACH Delaney (grant UM1 AI164565 to M. Caskey and R. B. Jones), the Einstein-Rockefeller-CUNY Center for AIDS Research (grant 1P30AI124414-01A1), BEAT-HIV Delaney (grant UM1 AI126620 to M. Caskey), the Bill and Melinda Gates Foundation (Collaboration for AIDS Vaccine discovery grant INV-002705) and the Stavros Niarchos Foundation (SNF) as part of its grant to the SNF Institute for Global Infectious Disease Research at The Rockefeller University. M.C.Nussenzweig is a Howard Hughes Medical Institute (HHMI) Investigator. This article is subject to HHMI’s Open Access to Publications policy. HHMI lab heads have previously granted a non-exclusive CC BY 4.0 license to the public and a sublicensable license to HHMI in their research articles. Pursuant to those licenses, the author-accepted manuscript of this article can be made freely available under a CC BY 4.0 license immediately upon publication. The authors declare no competing financial interests.

## References

Amemiya, H.M., A. Kundaje, and A.P. Boyle. 2019. The ENCODE Blacklist: Identification of Problematic Regions of the Genome. Sci Rep 9:9354.

Antar, A.A.R., K.M. Jenike, S. Jang, D.N. Rigau, D.B. Reeves, R. Hoh, M.R. Krone, J.C. Keruly, R.D. Moore, J.T. Schiffer, B.A.S. Nonyane, F.M. Hecht, S.G. Deeks, J.D. Siliciano, Y.-C. Ho, and R.F. Siliciano. 2020. Longitudinal study reveals HIV-1– infected CD4+ T cell dynamics during long-term antiretroviral therapy. Journal of Clinical Investigation 130:3543–3559.

Bachmann, N., C. von Siebenthal, V. Vongrad, T. Turk, K. Neumann, N. Beerenwinkel, J. Bogojeska, J. Fellay, V. Roth, Y.L. Kok, C.W. Thorball, A. Borghesi, S. Parbhoo, M. Wieser, J. Boni, M. Perreau, T. Klimkait, S. Yerly, M. Battegay, A. Rauch, M. Hoffmann, E. Bernasconi, M. Cavassini, R.D. Kouyos, H.F. Gunthard, K.J. Metzner, and H.I.V.C.S. Swiss. 2019. Determinants of HIV-1 reservoir size and long-term dynamics during suppressive ART. Nat Commun 10:3193.

Battivelli, E., M.S. Dahabieh, M. Abdel-Mohsen, J.P. Svensson, I. Tojal Da Silva, L.B. Cohn, A. Gramatica, S. Deeks, W.C. Greene, S.K. Pillai, and E. Verdin. 2018. Distinct chromatin functional states correlate with HIV latency reactivation in infected primary CD4(+) T cells. Elife 7:

Bui, J.K., M.D. Sobolewski, B.F. Keele, J. Spindler, A. Musick, A. Wiegand, B.T. Luke, W. Shao, S.H. Hughes, J.M. Coffin, M.F. Kearney, and J.W. Mellors. 2017. Proviruses with identical sequences comprise a large fraction of the replication-competent HIV reservoir. PLOS Pathogens 13:e1006283.

Cesana, D., F.R. Santoni de Sio, L. Rudilosso, P. Gallina, A. Calabria, S. Beretta, I. Merelli, E. Bruzzesi, L. Passerini, S. Nozza, E. Vicenzi, G. Poli, S. Gregori, G. Tambussi, and E. Montini. 2017. HIV-1-mediated insertional activation of STAT5B and BACH2 trigger viral reservoir in T regulatory cells. Nature Communications 8:498.

Chen, H.C., J.P. Martinez, E. Zorita, A. Meyerhans, and G.J. Filion. 2017. Position effects influence HIV latency reversal. Nat Struct Mol Biol 24:47–54.

Cho, A., C. Gaebler, T. Olveira, V. Ramos, M. Saad, J.C.C. Lorenzi, A. Gazumyan, S. Moir, M. Caskey, T.-W. Chun, and M.C. Nussenzweig. 2022. Longitudinal clonal dynamics of HIV-1 latent reservoirs measured by combination quadruplex polymerase chain reaction and sequencing. Proceedings of the National Academy of Sciences 119:e2117630119.

Chomont, N., M. El-Far, P. Ancuta, L. Trautmann, F.A. Procopio, B. Yassine-Diab, G. Boucher, M.-R. Boulassel, G. Ghattas, J.M. Brenchley, T.W. Schacker, B.J. Hill, D.C. Douek, J.-P. Routy, E.K. Haddad, and R.-P. Sékaly. 2009. HIV reservoir size and persistence are driven by T cell survival and homeostatic proliferation. Nature Medicine 15:893–900.

Chun, T.-W., D. Engel, M.M. Berrey, T. Shea, L. Corey, and A.S. Fauci. 1998. Early establishment of a pool of latently infected, resting CD4 ^+^ T cells during primary HIV-1 infection. Proceedings of the National Academy of Sciences 95:8869–8873.

Cohn, L.B., N. Chomont, and S.G. Deeks. 2020. The Biology of the HIV-1 Latent Reservoir and Implications for Cure Strategies. Cell Host & Microbe 27:519–530.

Cohn, L.B., I.T. da Silva, R. Valieris, A.S. Huang, J.C.C. Lorenzi, Y.Z. Cohen, J.A. Pai, A.L. Butler, M. Caskey, M. Jankovic, and M.C. Nussenzweig. 2018. Clonal CD4+ T cells in the HIV-1 latent reservoir display a distinct gene profile upon reactivation. Nature Medicine 24:604–609.

Cohn, L.B., I.T. Silva, T.Y. Oliveira, R.A. Rosales, E.H. Parrish, G.H. Learn, B.H. Hahn, J.L. Czartoski, M.J. McElrath, C. Lehmann, F. Klein, M. Caskey, B.D. Walker, J.D. Siliciano, R.F. Siliciano, M. Jankovic, and M.C. Nussenzweig. 2015. HIV-1 integration landscape during latent and active infection. Cell 160:420–432.

Collora, J.A., and Y.C. Ho. 2023. Integration site-dependent HIV-1 promoter activity shapes host chromatin conformation. Genome Res 33:891–906.

Collora, J.A., R. Liu, D. Pinto-Santini, N. Ravindra, C. Ganoza, J.R. Lama, R. Alfaro, J. Chiarella, S. Spudich, K. Mounzer, P. Tebas, L.J. Montaner, D. van Dijk, A. Duerr, and Y.-C. Ho. 2022. Single-cell multiomics reveals persistence of HIV-1 in expanded cytotoxic T cell clones. Immunity 55:1013–1031.e1017.

Craigie, R., and F.D. Bushman. 2012. HIV DNA Integration. Cold Spring Harbor Perspectives in Medicine 2:a006890.

Crooks, A.M., R. Bateson, A.B. Cope, N.P. Dahl, M.K. Griggs, J.D. Kuruc, C.L. Gay, J.J. Eron, D.M. Margolis, R.J. Bosch, and N.M. Archin. 2015. Precise Quantitation of the Latent HIV-1 Reservoir: Implications for Eradication Strategies. The Journal of Infectious Diseases 212:1361–1365.

Darcis, G., B. Berkhout, and A.O. Pasternak. 2019. The quest for cellular markers of HIV reservoirs: Any color you like. Frontiers in Immunology 10:

Demoustier, A., B. Gubler, O. Lambotte, M.G. de Goer, C. Wallon, C. Goujard, J.F. Delfraissy, and Y. Taoufik. 2002. In participants on prolonged HAART, a significant pool of HIV infected CD4 T cells are HIV-specific. AIDS 16:1749–1754.

Douek, D.C., J.M. Brenchley, M.R. Betts, D.R. Ambrozak, B.J. Hill, Y. Okamoto, J.P. Casazza, J. Kuruppu, K. Kunstman, S. Wolinsky, Z. Grossman, M. Dybul, A. Oxenius, D.A. Price, M. Connors, and R.A. Koup. 2002. HIV preferentially infects HIV-specific CD4+ T cells. Nature 417:95–98.

Dufour, C., P. Gantner, R. Fromentin, and N. Chomont. 2021. The multifaceted nature of HIV latency. In.

Einkauf, K.B., G.Q. Lee, C. Gao, R. Sharaf, X. Sun, S. Hua, S.M.Y. Chen, C. Jiang, X. Lian, F.Z. Chowdhury, E.S. Rosenberg, T.-W. Chun, J.Z. Li, X.G. Yu, and M. Lichterfeld. 2019. Intact HIV-1 proviruses accumulate at distinct chromosomal positions during prolonged antiretroviral therapy. Journal of Clinical Investigation 129:988–998.

Einkauf, K.B., M.R. Osborn, C. Gao, W. Sun, X. Sun, X. Lian, E.M. Parsons, G.T. Gladkov, K.W. Seiger, J.E. Blackmer, C. Jiang, S.A. Yukl, E.S. Rosenberg, X.G. Yu, and M. Lichterfeld. 2022. Parallel analysis of transcription, integration, and sequence of single HIV-1 proviruses. Cell 185:266–282 e215.

Finzi, D., J. Blankson, J.D. Siliciano, J.B. Margolick, K. Chadwick, T. Pierson, K. Smith, J. Lisziewicz, F. Lori, C. Flexner, T.C. Quinn, R.E. Chaisson, E. Rosenberg, B. Walker, S. Gange, J. Gallant, and R.F. Siliciano. 1999. Latent infection of CD4+ T cells provides a mechanism for lifelong persistence of HIV-1, even in participants on effective combination therapy. Nat Med 5:512–517.

Gaebler, C., J.C.C. Lorenzi, T.Y. Oliveira, L. Nogueira, V. Ramos, C.L. Lu, J.A. Pai, P. Mendoza, M. Jankovic, M. Caskey, and M.C. Nussenzweig. 2019. Combination of quadruplex qPCR and next-generation sequencing for qualitative and quantitative analysis of the HIV-1 latent reservoir. J Exp Med 216:2253–2264.

Gibson, D.G., L. Young, R.-Y. Chuang, J.C. Venter, C.A.H. 3rd, and H.O. Smith. 2009. Enzymatic assembly of DNA molecules up to several hundred kilobases. Nature Methods 6:343–345.

Han, Y., K. Lassen, D. Monie, A.R. Sedaghat, S. Shimoji, X. Liu, T.C. Pierson, J.B. Margolick, R.F. Siliciano, and J.D. Siliciano. 2004. Resting CD4+ T cells from human immunodeficiency virus type 1 (HIV-1)-infected individuals carry integrated HIV-1 genomes within actively transcribed host genes. J Virol 78:6122–6133.

Hartweger, H., R. Gautam, Y. Nishimura, F. Schmidt, K.-H. Yao, A. Escolano, M. Jankovic, M.A. Martin, and M.C. Nussenzweig. 2023. Gene Editing of Primary Rhesus Macaque B Cells. J. Vis. Exp 192:

Ho, Y.-C., L. Shan, Nina N. Hosmane, J. Wang, Sarah B. Laskey, Daniel I.S. Rosenbloom, J. Lai, Joel N. Blankson, Janet D. Siliciano, and Robert F. Siliciano. 2013. Replication-Competent Noninduced Proviruses in the Latent Reservoir Increase Barrier to HIV-1 Cure. Cell 155:540–551.

Hosmane, N.N., K.J. Kwon, K.M. Bruner, A.A. Capoferri, S. Beg, D.I.S. Rosenbloom, B.F. Keele, Y.-C. Ho, J.D. Siliciano, and R.F. Siliciano. 2017. Proliferation of latently infected CD4+ T cells carrying replication-competent HIV-1: Potential role in latent reservoir dynamics. Journal of Experimental Medicine 214:959–972.

Huang, A.S., V. Ramos, T.Y. Oliveira, C. Gaebler, M. Jankovic, M.C. Nussenzweig, and L.B. Cohn. 2021. Integration features of intact latent HIV-1 in CD4+ T cell clones contribute to viral persistence. J Exp Med 218:

Hultquist, J.F., J. Hiatt, K. Schumann, M.J. McGregor, T.L. Roth, P. Haas, J.A. Doudna, A. Marson, and N.J. Krogan. 2019. CRISPR-Cas9 genome engineering of primary CD4(+) T cells for the interrogation of HIV-host factor interactions. Nat Protoc 14:1–27.

Jiang, C., X. Lian, C. Gao, X. Sun, K.B. Einkauf, J.M. Chevalier, S.M.Y. Chen, S. Hua, B. Rhee, K. Chang, J.E. Blackmer, M. Osborn, M.J. Peluso, R. Hoh, M. Somsouk, J. Milush, L.N. Bertagnolli, S.E. Sweet, J.A. Varriale, P.D. Burbelo, T.W. Chun, G.M. Laird, E. Serrao, A.N. Engelman, M. Carrington, R.F. Siliciano, J.M. Siliciano, S.G. Deeks, B.D. Walker, M. Lichterfeld, and X.G. Yu. 2020. Distinct viral reservoirs in individuals with spontaneous control of HIV-1. Nature 585:261–267.

Jones, R.B., C. Kovacs, T.W. Chun, and M.A. Ostrowski. 2012. Short communication: HIV type 1 accumulates in influenza-specific T cells in subjects receiving seasonal vaccination in the context of effective antiretroviral therapy. AIDS Res Hum Retroviruses 28:1687–1692.

Jones, R.B., S. Mueller, R. O’Connor, K. Rimpel, D.D. Sloan, D. Karel, H.C. Wong, E.K. Jeng, A.S. Thomas, J.B. Whitney, S.-Y. Lim, C. Kovacs, E. Benko, S. Karandish, S.-H. Huang, M.J. Buzon, M. Lichterfeld, A. Irrinki, J.P. Murry, A. Tsai, H. Yu, R. Geleziunas, A. Trocha, M.A. Ostrowski, D.J. Irvine, and B.D. Walker. 2016. A Subset of Latency-Reversing Agents Expose HIV-Infected Resting CD4+ T-Cells to Recognition by Cytotoxic T-Lymphocytes. PLOS Pathogens 12:e1005545.

Jordan, A. 2003. HIV reproducibly establishes a latent infection after acute infection of T cells in vitro. The EMBO Journal 22:1868–1877.

Jordan, A., P. Defechereux, and E. Verdin. 2001. The site of HIV-1 integration in the human genome determines basal transcriptional activity and response to Tat transactivation. The EMBO Journal 20:1726–1738.

Lee, G.Q., N. Orlova-Fink, K. Einkauf, F.Z. Chowdhury, X. Sun, S. Harrington, H.H. Kuo, S. Hua, H.R. Chen, Z. Ouyang, K. Reddy, K. Dong, T. Ndung’u, B.D. Walker, E.S. Rosenberg, X.G. Yu, and M. Lichterfeld. 2017. Clonal expansion of genome-intact HIV-1 in functionally polarized Th1 CD4+ T cells. J Clin Invest 127:2689–2696.

Lewinski, M.K., D. Bisgrove, P. Shinn, H. Chen, C. Hoffmann, S. Hannenhalli, E. Verdin, C.C. Berry, J.R. Ecker, and F.D. Bushman. 2005. Genome-wide analysis of chromosomal features repressing human immunodeficiency virus transcription. J Virol 79:6610–6619.

Li, J.Z., B. Etemad, H. Ahmed, E. Aga, R.J. Bosch, J.W. Mellors, D.R. Kuritzkes, M.M. Lederman, M. Para, and R.T. Gandhi. 2016. The size of the expressed HIV reservoir predicts timing of viral rebound after treatment interruption. *AIDS (London*, England) 30:343–353.

Lichterfeld, M., C. Gao, and X.G. Yu. 2022. An ordeal that does not heal: understanding barriers to a cure for HIV-1 infection. Trends in Immunology 43:608–616.

Liu, R., F.R. Simonetti, and Y.-C. Ho. 2020. The forces driving clonal expansion of the HIV-1 latent reservoir. Virology Journal 17:4.

Lorenzi, J.C.C., Y.Z. Cohen, L.B. Cohn, E.F. Kreider, J.P. Barton, G.H. Learn, T. Oliveira, C.L. Lavine, J.A. Horwitz, A. Settler, M. Jankovic, M.S. Seaman, A.K. Chakraborty, B.H. Hahn, M. Caskey, and M.C. Nussenzweig. 2016. Paired quantitative and qualitative assessment of the replication-competent HIV-1 reservoir and comparison with integrated proviral DNA. Proceedings of the National Academy of Sciences 113:E7908–E7916.

Lu, C.-L., J.A. Pai, L. Nogueira, P. Mendoza, H. Gruell, T.Y. Oliveira, J. Barton, J.C.C. Lorenzi, Y.Z. Cohen, L.B. Cohn, F. Klein, M. Caskey, M.C. Nussenzweig, and M. Jankovic. 2018. Relationship between intact HIV-1 proviruses in circulating CD4+ T cells and rebound viruses emerging during treatment interruption. Proceedings of the National Academy of Sciences 115:E11341–E11348.

Malim, M.H., and M. Emerman. 2008. HIV-1 Accessory Proteins—Ensuring Viral Survival in a Hostile Environment. Cell Host & Microbe 3:388–398.

Margolis, D.M., and N.M. Archin. 2017. Proviral Latency, Persistent Human Immunodeficiency Virus Infection, and the Development of Latency Reversing Agents. The Journal of Infectious Diseases 215:S111–S118.

Martin, R.M., K. Ikeda, M.K. Cromer, N. Uchida, T. Nishimura, R. Romano, A.J. Tong, V.T. Lemgart, J. Camarena, M. Pavel-Dinu, C. Sindhu, V. Wiebking, S. Vaidyanathan, D.P. Dever, R.O. Bak, A. Laustsen, B.J. Lesch, M.R. Jakobsen, V. Sebastiano, H. Nakauchi, and M.H. Porteus. 2019. Highly Efficient and Marker-free Genome Editing of Human Pluripotent Stem Cells by CRISPR-Cas9 RNP and AAV6 Donor-Mediated Homologous Recombination. Cell Stem Cell 24:821–828 e825.

Mendoza, P., J.R. Jackson, T.Y. Oliveira, C. Gaebler, V. Ramos, M. Caskey, M. Jankovic, M.C. Nussenzweig, and L.B. Cohn. 2020. Antigen-responsive CD4+ T cell clones contribute to the HIV-1 latent reservoir. J Exp Med 217:

Mori, L., and S.T. Valente. 2020. Key Players in HIV-1 Transcriptional Regulation: Targets for a Functional Cure. Viruses 12:

Ne, E., R.-J. Palstra, and T. Mahmoudi. 2018. Chapter Six - Transcription: Insights From the HIV-1 Promoter. In International Review of Cell and Molecular Biology. F. Loos, editor Academic Press, 191–243.

Pace, M.J., L. Agosto, E.H. Graf, and U. O’Doherty. 2011. HIV reservoirs and latency models. Virology 411:344–354.

Palmer, S., A.P. Wiegand, F. Maldarelli, H. Bazmi, J.M. Mican, M. Polis, R.L. Dewar, A. Planta, S. Liu, J.A. Metcalf, J.W. Mellors, and J.M. Coffin. 2003. New real-time reverse transcriptase-initiated PCR assay with single-copy sensitivity for human immunodeficiency virus type 1 RNA in plasma. J Clin Microbiol 41:4531–4536.

Peluso, M.J., P. Bacchetti, K.D. Ritter, S. Beg, J. Lai, J.N. Martin, P.W. Hunt, T.J. Henrich, J.D. Siliciano, R.F. Siliciano, G.M. Laird, and S.G. Deeks. 2020. Differential decay of intact and defective proviral DNA in HIV-1–infected individuals on suppressive antiretroviral therapy. JCI Insight 5:

Perelson, A.S., A.U. Neumann, M. Markowitz, J.M. Leonard, and D.D. Ho. 1996. HIV-1 dynamics in vivo: virion clearance rate, infected cell life-span, and viral generation time. Science 271:1582–1586.

Pinzone, M.R., D.J. VanBelzen, S. Weissman, M.P. Bertuccio, L. Cannon, E. Venanzi-Rullo, S. Migueles, R.B. Jones, T. Mota, S.B. Joseph, K. Groen, A.O. Pasternak, W.-T. Hwang, B. Sherman, A. Vourekas, G. Nunnari, and U. O’Doherty. 2019. Longitudinal HIV sequencing reveals reservoir expression leading to decay which is obscured by clonal expansion. Nature Communications 10:728.

Raj, A., and A.v. Oudenaarden. 2008. Nature, Nurture, or Chance: Stochastic Gene Expression and Its Consequences. Cell 135:216–226.

Salazar-Gonzalez, J.F., E. Bailes, K.T. Pham, M.G. Salazar, M.B. Guffey, B.F. Keele, C.A. Derdeyn, P. Farmer, E. Hunter, S. Allen, O. Manigart, J. Mulenga, J.A. Anderson, R. Swanstrom, B.F. Haynes, G.S. Athreya, B.T. Korber, P.M. Sharp, G.M. Shaw, and B.H. Hahn. 2008. Deciphering human immunodeficiency virus type 1 transmission and early envelope diversification by single-genome amplification and sequencing. J Virol 82:3952–3970.

Saleh, S., F. Wightman, S. Ramanayake, M. Alexander, N. Kumar, G. Khoury, C. Pereira, D. Purcell, P.U. Cameron, and S.R. Lewin. 2011. Expression and reactivation of HIV in a chemokine induced model of HIV latency in primary resting CD4+ T cells. Retrovirology 8:80.

Scheerder, M.-A.D., B. Vrancken, S. Dellicour, T. Schlub, E. Lee, W. Shao, S. Rutsaert, C. Verhofstede, T. Kerre, T. Malfait, D. Hemelsoet, M. Coppens, A. Dhondt, D.D. Looze, F. Vermassen, P. Lemey, S. Palmer, and L. Vandekerckhove. 2019. HIV Rebound Is Predominantly Fueled by Genetically Identical Viral Expansions from Diverse Reservoirs. Cell Host & Microbe 26:347–358.e347.

Schröder, A.R.W., P. Shinn, H. Chen, C. Berry, J.R. Ecker, and F. Bushman. 2002. HIV-1 Integration in the Human Genome Favors Active Genes and Local Hotspots. Cell 110:521–529.

Sherrill-Mix, S., M.K. Lewinski, M. Famiglietti, A. Bosque, N. Malani, K.E. Ocwieja, C.C. Berry, D. Looney, L. Shan, L.M. Agosto, M.J. Pace, R.F. Siliciano, U. O’Doherty, J. Guatelli, V. Planelles, and F.D. Bushman. 2013. HIV latency and integration site placement in five cell-based models. Retrovirology 10:90.

Siliciano, J.D., J. Kajdas, D. Finzi, T.C. Quinn, K. Chadwick, J.B. Margolick, C. Kovacs, S.J. Gange, and R.F. Siliciano. 2003. Long-term follow-up studies confirm the stability of the latent reservoir for HIV-1 in resting CD4+ T cells. Nat Med 9:727–728.

Siliciano, J.D., and R.F. Siliciano. 2022. In Vivo Dynamics of the Latent Reservoir for HIV-1: New Insights and Implications for Cure. Annual Review of Pathology: Mechanisms of Disease 17:271–294.

Simonetti, F.R., M.D. Sobolewski, E. Fyne, W. Shao, J. Spindler, J. Hattori, E.M. Anderson, S.A. Watters, S. Hill, X. Wu, D. Wells, L. Su, B.T. Luke, E.K. Halvas, G. Besson, K.J. Penrose, Z. Yang, R.W. Kwan, C. Van Waes, T. Uldrick, D.E. Citrin, J. Kovacs, M.A. Polis, C.A. Rehm, R. Gorelick, M. Piatak, B.F. Keele, M.F. Kearney, J.M. Coffin, S.H. Hughes, J.W. Mellors, and F. Maldarelli. 2016. Clonally expanded CD4+ T cells can produce infectious HIV-1 in vivo. Proceedings of the National Academy of Sciences of the United States of America 113:1883–1888.

Simonetti, F.R., H. Zhang, G.P. Soroosh, J. Duan, K. Rhodehouse, A.L. Hill, S.A. Beg, K. McCormick, H.E. Raymond, C.L. Nobles, J.K. Everett, K.J. Kwon, J.A. White, J. Lai, J.B. Margolick, R. Hoh, S.G. Deeks, F.D. Bushman, J.D. Siliciano, and R.F. Siliciano. 2021. Antigen-driven clonal selection shapes the persistence of HIV-1–infected CD4^+^ T cells in vivo. The Journal of Clinical Investigation 131:

Skupsky, R., J.C. Burnett, J.E. Foley, D.V. Schaffer, and A.P. Arkin. 2010. HIV promoter integration site primarily modulates transcriptional burst size rather than frequency. PLoS Comput Biol 6:

Sun, W., C. Gao, C.A. Hartana, M.R. Osborn, K.B. Einkauf, X. Lian, B. Bone, N. Bonheur, T.W. Chun, E.S. Rosenberg, B.D. Walker, X.G. Yu, and M. Lichterfeld. 2023. Phenotypic signatures of immune selection in HIV-1 reservoir cells. Nature 614:309–317.

Wei, X., S.K. Ghosh, M.E. Taylor, V.A. Johnson, E.A. Emini, P. Deutsch, J.D. Lifson, S. Bonhoeffer, M.A. Nowak, B.H. Hahn, and, et al. 1995. Viral dynamics in human immunodeficiency virus type 1 infection. Nature 373:117–122.

Weymar, G.H.J., Y. Bar-On, T.Y. Oliveira, C. Gaebler, V. Ramos, H. Hartweger, G. Breton, M. Caskey, L.B. Cohn, M. Jankovic, and M.C. Nussenzweig. 2022. Distinct gene expression by expanded clones of quiescent memory CD4(+) T cells harboring intact latent HIV-1 proviruses. Cell Rep 40:111311.

Wong, J.K., M. Hezareh, H.F. Günthard, D.V. Havlir, C.C. Ignacio, C.A. Spina, and D.D. Richman. 1997. Recovery of Replication-Competent HIV Despite Prolonged Suppression of Plasma Viremia. Science 278:1291–1295.

Wong, M., Y. Wei, and Y.-C. Ho. 2023. Single-cell multiomic understanding of HIV-1 reservoir at epigenetic, transcriptional, and protein levels. Current opinion in HIV and AIDS 18:246–256.

Wu, V.H., J.M.L. Nordin, S. Nguyen, J. Joy, F. Mampe, P.M. Del Rio Estrada, F. Torres-Ruiz, M. Gonzalez-Navarro, Y.A. Luna-Villalobos, S. Avila-Rios, G. Reyes-Teran, P. Tebas, L.J. Montaner, K.J. Bar, L.A. Vella, and M.R. Betts. 2023. Profound phenotypic and epigenetic heterogeneity of the HIV-1-infected CD4(+) T cell reservoir. Nat Immunol 24:359–370.

Yeh, Y.-H.J., K.M. Jenike, R.M. Calvi, J. Chiarella, R. Hoh, S.G. Deeks, and Y.-C. Ho. 2020. Filgotinib suppresses HIV-1–driven gene transcription by inhibiting HIV-1 splicing and T cell activation. The Journal of Clinical Investigation 130:4969–4984.

