## Supplementary material for "Transcription of HIV-1 at sites of intact latent provirus integration": Table S1 and Figures S1-5

**Table S1. Clinical characteristics of study participants**. Dx: diagnosis; ART: antiretroviral treatment; EFV: Efavirenz; TDF: tenofovir disoproxil fumarate; FTC: emtricitabine; DTG: dolutegravir.

| ID | Age | Sex | Race | Year HIV-1 Dx | Year ART initiation | Years of uninterrupted ART | Viral Load (copies/mL) | $CD4^+$ T cell count | Reported Nadir | ART Regimen |
|---|---|---|---|---|---|---|---|---|---|---|
| 603 | 43 | Male | White/ Hispanic | 12 | 10 | 10 | <20 | 300 | 693 | EFV/TDF /FTC |
| 5104 | 35 | Male | Black | 7 | 7 | 7 | <20 | 450 | 1006 | DTG/TD F/FTC |

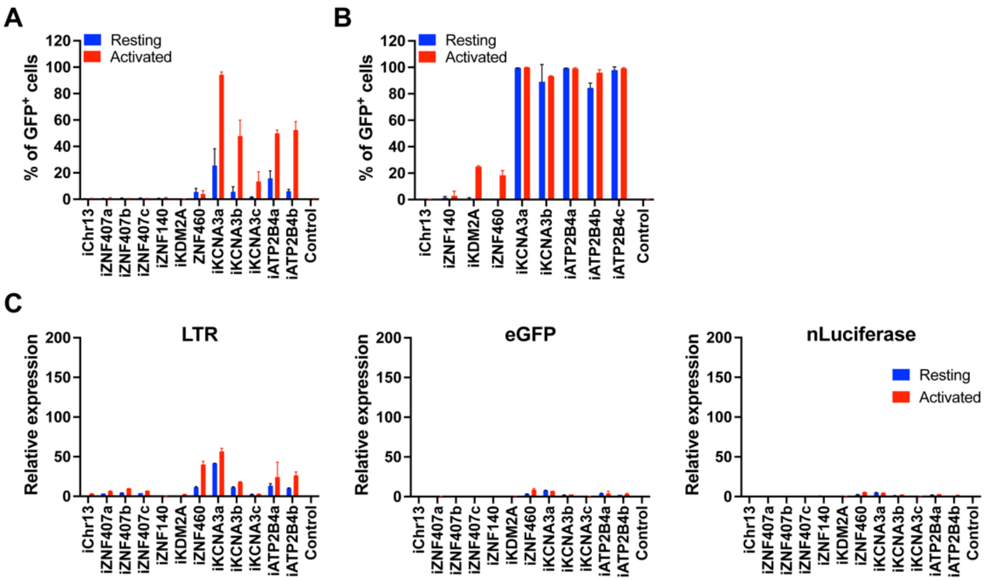


**Figure S1. GFP expression in reporter cell lines. (A-B)** Percentage of GFP+ cells as measured by flow cytometry for each integration-positive clone and control, under resting (blue) and PMA/ionomycin- and CD3/CD28-activated (red) conditions, respectively for Jurkat **(A)** and primary CD4$^{+}$ T **(B)** cells. Bars represent the mean of two independent experiments (biological replicates) ± standard deviation. **(C)** Graphs showing relative LTR (left panel), eGFP (middle panel) and nLuciferase (right panel) expression assessed by qPCR in Jurkat cell clones under resting (blue) and CD3/CD28-activated (red) conditions. Bars represent the mean relative expression from two independent assays (biological replicates) ± standard deviation.

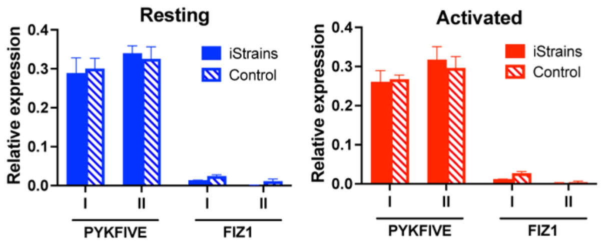


**Figure S2. Expression host genes carrying reporter proviruses in Jurkat clones.** Relative expression of host genes *PYKFYVE* and *Fiz1* in Jurkat clones (full bars) and control (stripped bars), under resting (blue) and activated (red) conditions. Bars represent the mean relative expression ± standard deviation for each gene for three technical replicates, for two independent assays (biological replicates**,** I and II).

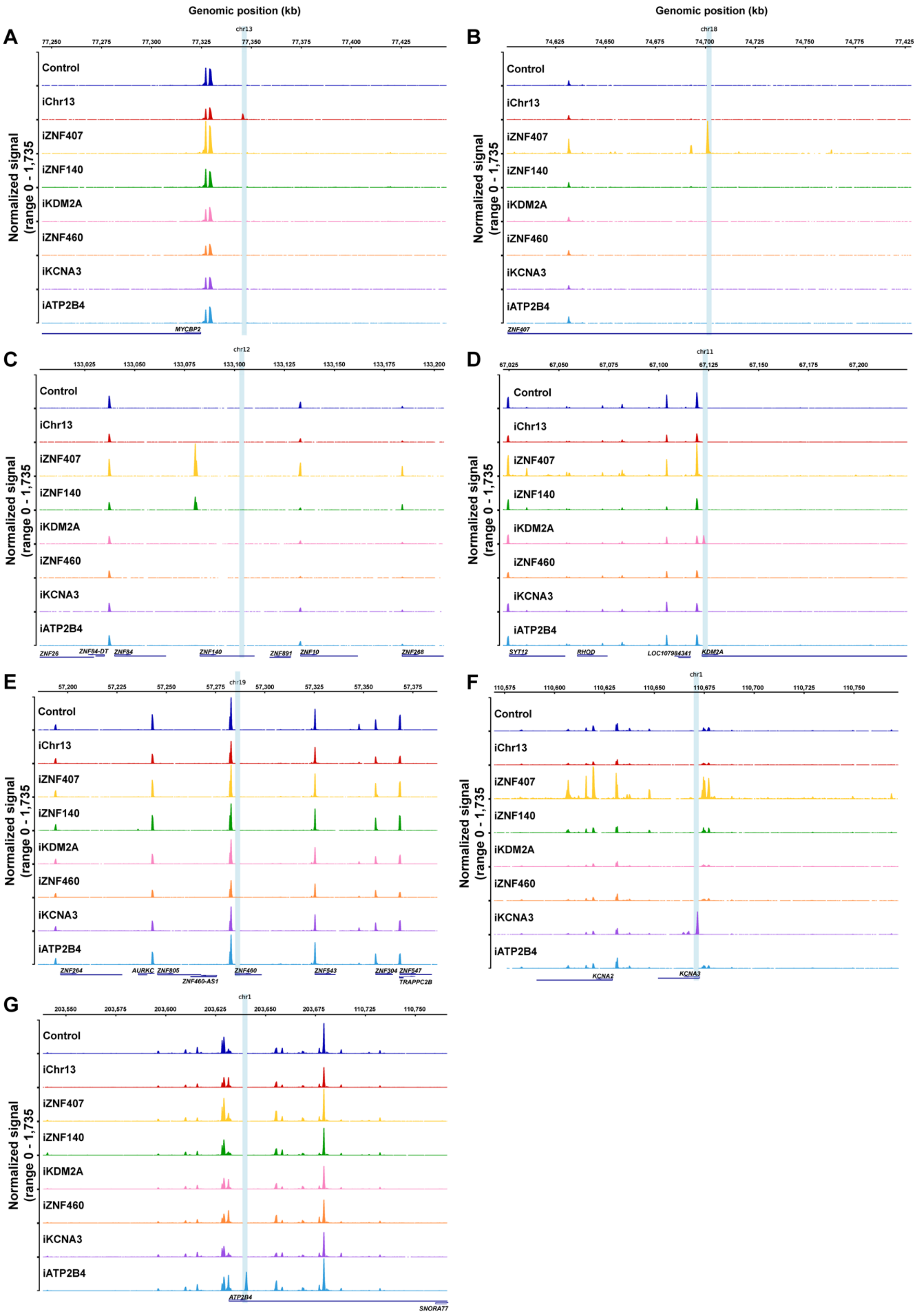
Genomic position (kb)
Genomic position (kb)
A
chr13
77,250
77,275
77,300
77,325
77,350
77,375
77,400
77,425
Normalized signal
(range 0 - 1,735
Control
iChr13
iZNF407
iZNF140
iKDM2A
iZNF460
iKCNA3
iATP2B4
MYCBP2
B
chr18
74,625
74,650
74,675
74,700
74,725
74,750
74,775
77,425
ZNF407
C
chr12
133,025
133,050
133,075
133,100
133,125
133,150
133,175
133,200
ZNF26
ZNF84-DT
ZNF84
ZNF140
ZNF891
ZNF10
ZNF268
D
chr11
67,025
67,050
67,075
67,100
67,125
67,150
67,175
67,200
SYT12
RHOD
LOC107984341
KDM2A
E
chr19
57,200
57,225
57,250
57,275
57,300
57,325
57,350
57,375
ZNF264
AURKC
ZNF805
ZNF460-AS1
ZNF460
ZNF543
ZNF304
ZNF547
TRAPPC2B
F
chr1
110,575
110,600
110,625
110,650
110,675
110,700
110,725
110,750
KCNA2
KCNA3
G
chr1
203,550
203,575
203,600
203,625
203,650
203,675
110,725
110,750
ATP2B4
SNORA77

**Figure S3. Chromatin accessibility around reporter construct integration site in Jurkat cell clones. (A-G)** Chromatin accessibility measured by ATAC-Seq in a 200,000 kb window of the genome around each of the integration sites for all Jurkat clones: chr13 (**A**), chr18 (*ZNF407*, **B**), chr12 (*ZNF140*, **C**), chr11 (*KDM2A*, **D**), chr19 (*ZNF460*, **E**), chr1 (*KCNA3*, **F**) and chr1 (*ATP2B4*, **G**). Graphs were generated by averaging the normalized reads from three technical replicates for each clone.

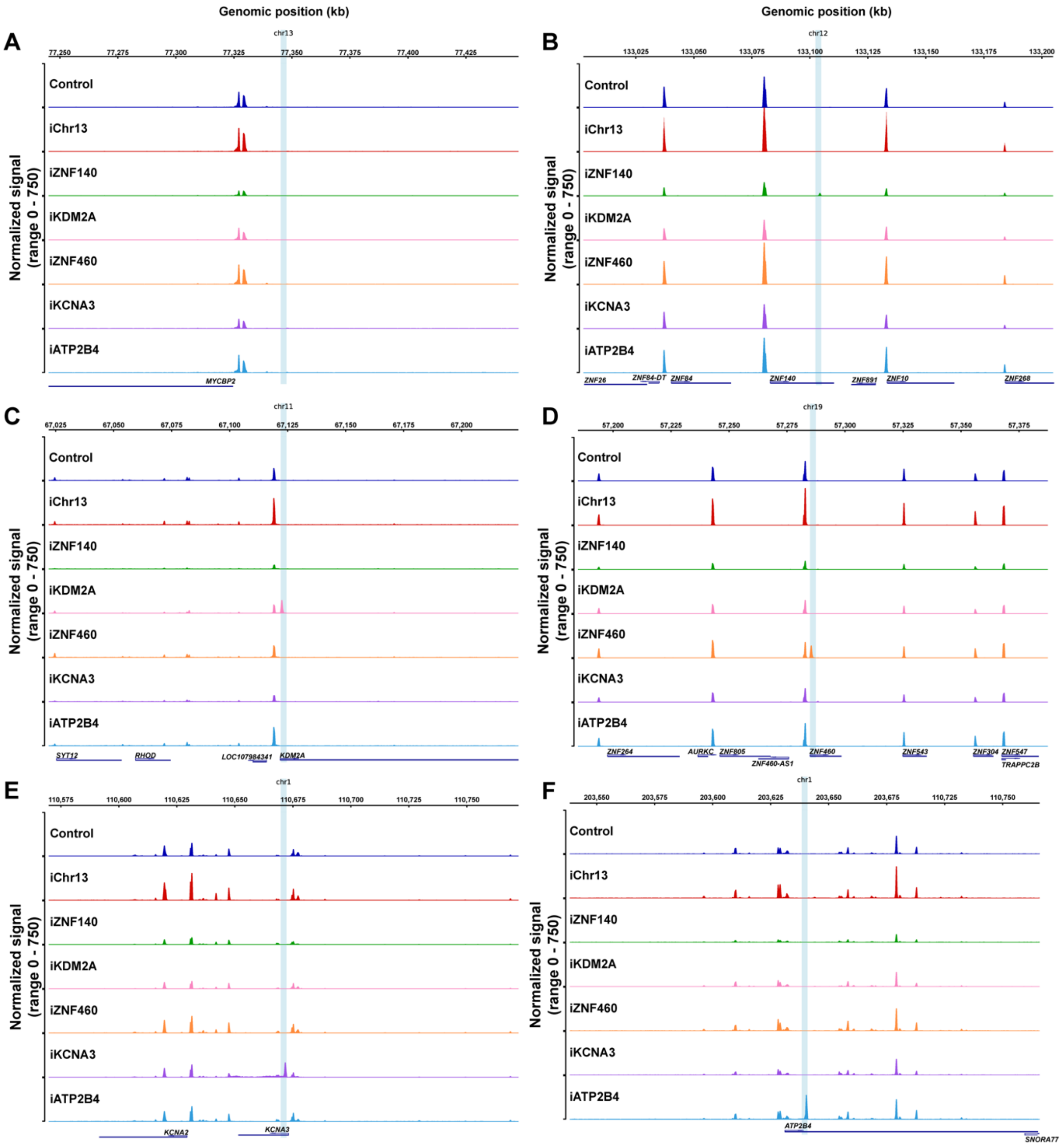


**Figure S4. Chromatin accessibility around reporter construct integration site in primary $CD4^+$ T cell clones. (A-F)** Chromatin accessibility measured by ATAC-Seq in a 200,000 kb window of the genome around each of the integration sites for all primary $CD4^+$ T cell clones: chr13 (**A**), chr12 (*ZNF140*, **B**), chr11 (*KDM2A*, **C**), chr19 (*ZNF460*, **D**), chr1 (*KCNA3*, **E**) and chr1 (*ATP2B4*, **F**). Graphs were generated by averaging the normalized reads from three technical replicates for each clone.

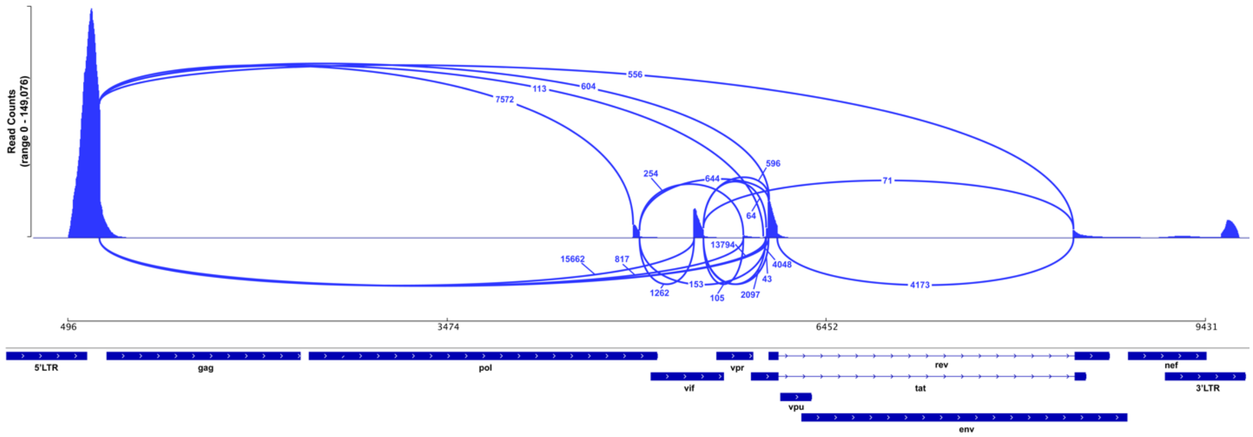


**Figure S5. HIV-1 RNA splice variants.** Sashimi plot showing scRNA-Seq reads of 10X single cell gene-expression data mapping to the HIV genome (bottom), in infected cells of participant 5104 under resting conditions. Splice junctions are represented as arcs connecting exons and the histogram represents the read coverage at each junction. The number of reads mapping to each junction is indicated by the numbers associated with each arc.
